# *Croton caudatus* Geiseler Induces Differentiation of the Acute Myeloid Leukemia cell lines

**DOI:** 10.1101/2022.06.30.498361

**Authors:** Takhellambam Chanu Machathoibi, Lisam Shanjukumar Singh

**Affiliations:** Recobinant DNA Technology, Medical Biotechnogy and Animal Behavior Lab, Department of Biotechnology, Manipur University, Canchipur, Imphal-795003, Manipur, India; Cancer and Molecular Biology Division, Department of Biotechnology, Manipur University, Canchipur, Imphal-795003, Manipur, India

**Keywords:** *Croton caudatus* Geiseler, HL60 differentiation, differentiation induction chemotherapy (DIC), Acute myeloid leukemia

## Abstract

*Croton caudatus* Geiseler (CCG) is a traditional medicinal plant and it has also been reported as having anticancer activity. However, most of the studies of the anticancer activity of CGG based only on cytotoxicity and antioxidant properties. The present study investigates the anticancer activity of leaf crude extract of CCG on an acute myeloid leukemia cell line, HL60. The results showed that CCG promotes the monocyte macrophage-like morphology of HL60 cells but it does not cause cytotoxicity on HL60.The results also showed that CCG increased the expression of a marker for monocyte/macrophage differentiation, CD11b, and induced arrest of cell cycle at S phase. Further, the results indicate that leaf the crude extract of CCG has a very high ROS activity. The present study is the first report of CCG inducing cell differentiation. The overall results suggest that CCG is a promising agent for differentiation induction chemotherapy (DIC) for leukemia.

## INTRODUCTION

Acute myeloid leukemia (AML) represents a heterogeneous malignancy characterized by clonal proliferation and impaired differentiation of myeloid precursors with diverse outcomes. Despite the advances in understanding the molecular heterogeneity and pathogenesis of AML, there has been little progress in the standard therapy for AML over the past four decades (1). It is the most common type of acute leukemia in adults (2). The American Cancer Society predicts that by 2020 there will be an estimated 21,040 new AML patients in the United States, and 11,180 people will die from this disease (3). AML accounts for 15% to 20% of all childhood leukemia (4). However, a reliable figure for the incidence of AML in Indian children is lacking (5). A characteristic abnormality of leukemia cells is that cells are blocked at an early stage of their development and therefore, fail to differentiate into functional mature cells. During the 1970s and 1980s, it came to an understanding that inducing malignant cells to overcome the blockage of differentiation and then enter the apoptotic pathways as an alternative to kill cancer cells by cytotoxic therapies (6). Differentiation induction chemotherapy (DIC), using chemical agents such as 1,25-dihydroxy vitamin D3 (D3), all trans retinoic acid (ATRA), retinoids, is a promising approach for the treatment of many cancers. The paradigm that leukemias are characterized by the alteration of sets of 2 genes; the genes that give the malignancy a proliferative advantage and the genes that are associated with blockage of cell differentiation, is still as valid today as it was three decades ago (6). The observations in the model systems, such as the lineage-uncommitted human myeloblastic cell line HL-60 that closely resemble patient-derived cells, could be used for understanding the differentiation programs occurring in patients. HL-60 undergoes cell cycle arrest and either myeloid or monocytic differentiation following stimulation; ATRA induces G1/G0-arrest and myeloid differentiation in HL-60 cell, while 1,25-dihydroxy vitamin D3 (D3) induces arrest and monocytic differentiation. Commitment to cell cycle arrest and differentiation requires approximately 48 hr of treatment, during which HL-60 cells undergo two division cycles. HL-60 cell line has been used as an established in-vitro model for studying various cellular and biochemical pathways involved in drug-induced differentiation.

*Croton caudatus* Geiseler (CCG) belongs to the Euphorbiaceae family (7). It is fairly widespread in South East Asia including Sri Lanka, Bhutan, Borneo, Burma, Indo-Myanmar region, Java, Laos, Malaysia, Nepal, Pakistan, Philippines, Singapore, Sumatra,Thailand, and Vietnam. Croton caudatus Geiseler had long been known to inhabitants of Manipur (8). It is traditionally used as a poultice for fever, sprains, and treatment of liver diseases in various parts of Asia (9). Its roots are purgative. It has been reported that the whole plant is used for medicinal purposes and it is found to be low in toxicity. *Croton caudatus* is a traditional Dai Nationalistic medicine of China. The stems and leaves of *Croton caudatus* have been used for the treatment of malaria, ardent fever, convulsions, rheumatic arthritis, and numbness (9,10). The leaves have been applied to festering wounds of injured cattle to ward off against maggots. Also, the use of *Croton caudatus* Geiseler in the treatment of cancer in the Saikot area of Manipur has been recently reported (11). However, studies on the anticancer activity of Croton caudatus Geiseler are rare. The present study investigates the anticancer activity of Croton caudatus Geiseler on HL60cell line.

## MATERIALS AND METHODS

### 1. Collection of Plant

The plant *Croton caudatus* Geiseler was collected from the Saikot village, Churachandpur district Manipur, about 65 km from Imphal at the latitude 24°3316595’N and longitude 93°7270017’E during the dry season. The cleaned and non-infected matured leaves were collected and washed with distilled water three times and dried in shade at room temperature in dried, clean, and hygienic conditions to avoid entry of insects, animals, fungus, and extraneous terrestrial material. The exhaust and free air circulation were allowed in the room. Gross particles like midribs were removed before processing. The leaves of the plant were grounded to a fine powder using an electronic grinder. The powder of the leaf thus prepared was stored at -20C in plastic bags till used.

### 2. Leaf extraction from *Croton caudatus* Geiseler

100 gm of the leaf powder was weighed and subjected to methanol (CH3OH, analytical grade) extraction using a soxhlet apparatus (500ml capacity) below 65°C. The crude extract obtained was evaporated in the air or using a lyophilizer. The extract was reconstituted in DMSO (Me_2_SO, cell culture grade) at a final stock concentration of 100μg/μl and filtered through a Whatmann syringe microfilter of pore size 0.2μm, made aliquotes into a small volume (20μl). The crude methanol extract (CME) of CCG thus prepared was stored at -20°C until further experiments.

### 3. Cell Culture

HL60 cells were obtained from the National Centre for Cell Science (NCCS) Pune, Maharashtra. Cells were grown in a culture flask with DMEM, supplemented with 10 % (v/v) FBS (Gibco) and 5% antibiotics penstrep (Gibco). The cells were all grown in optimal growth conditions of 37°C and 5% CO2 in a humidified chamber. The cells were passaged, and stocks were prepared and stored at liquid nitrogen. Cell lines were checked for any contamination and healthy condition under a microscope before each use. Only the uncontaminated and healthy cells were used. To avoid mycoplasma contamination, cells were treated with Plasmocin (InvivoGen) once in one to two months.

### 4. Cell Treatment with leave crude extract

About 2×105 HL60 cells were seeded in a 12-well culture plates and grown overnight in a cell culture incubator. The cells were treated with different doses of the leaf crude extract of CCG for different time points as mentioned elsewhere.

### 5. Geimsa staining

HL60 cells were grown for 24 hrs (70 to 75% confluency) and treated with 1μg/ml of leaf crude extract of CCG for 24, 48, and 72 hrs. Control cells were treated with DMSO (0.1% v/v), which was used as vehicle for extract treatment, in identical conditions. The media was removed, and the cells were washed with ice-cold 1XPBS. The cells were fixed with methanol for 10 mins, stained with Geimsa for 30 mins, and then the excess stain was removed with molecular grade water. The cells were observed in phase contrast microscope and pictures were captured in three different fields.

### 6. Analysis of CD11b expression by flowcytometry

HL60 cells were seeded at a confluency of 70% to 75% and after 24 hrs cells were treated with the 1μg/ml of leaf crude extract of CCG. Negative control cells were treated with DMSO and positive control cells were treated with 100nM of Phorbol 12-myristate 13-acetate (PMA) in identical conditions. About 5×105 HL60 cells were harvested and collected by centrifugation and washed twice with ice-cold 1X PBS, and the cells are resuspended in ice-cold PBS containing 10% FBS and 1% sodium azide, anti-human CD11b at a dilution of 1:200 (eBiosciences) was added and incubated for 30 mins at room temperature in the dark. The cells were washed with ice-cold PBS containing 10% FBS and 1% sodium azide for three times and incubated with 1μg/μl of rhodamine conjugated secondary antibody (Thermo Fisher Scientific) at a dilution of 1:200. The cells were washed three times with ice-cold PBS containing 10% FBS and 1% sodium azide. The quantitative immunoflorescence measurement was conducted by flow cytometry (Accrui 5, BD Biociences) and analyzed using Accrui 6, BD Bioscience software. At least 4×104 cells were collected and emission of the immunofloresence or rhodamine falls at FL2 channel i.e. at λ (wavelength) 535 nm (max). One-parameter histograms based upon the florescence of the cell were generated via the analysis of the least 4×104 cells recorded.

### 7. Cell cycle analysis by flowcytometer

HL60 cells were seeded in a 60 cm culture plate up to a confluence of 70% to 75%. HL60, cells were treated with 1μg/ml of leaf crude extract of CCG for 0hrs, 6 hrs, 12 hrs, 24 hrs, and 48hrs. Control cells were treated with DMSO. Media was removed and cells were washed with ice-cold 1XPBS, collected by trypsinization, washed twice with ice-cold 1XPBS. The cells were fixed with 70% ethanol, washed with 1XPBS two times, treated with DNase free RNase 100ng/ul (Quiagen) for 5min and then stained with fluorochrome solution containing 50 mg/ml of Propidium Iodide (PI), 0.1%Triton-x100, and 0.5% Sodium citrate. The quantitative fluorescence measurement was conducted on a flow cytometer (Accrui 5, BD Biociences) and analyzed using Accrui 6, BD Bioscience software. At least 1×104 cells were collected and emission of the fluorescence of PI falls at FL2 channel i.e. at λ (wavelength) 585 nm (max). One-parameter histogram based upon the florescence of the cell was generated via the analysis of the least 1X 104 cells recorded.

### 7. ROS detection by flowcytometry

HL60 cells were seeded at a confluency of 70% to 75% and after 24 hrs cells were treated with the 1μg/ml of leaf crude extract of CCG for 2 hrs. Negative control cells were treated with DMSO (0.1% v/v) and positive control cells were treated with 100μm H_2_O_2_ for 2 hrs in identical conditions. The cells were treated with 100nM DCFH-DA for 10 minutes in dark at 37C_J. Cells were harvested and resuspended in serum-free media without phenol red. About 1×104 cells were acquired per sample. The quantitative immunofluorescence measurement was conducted on a flow cytometer (Accrui 5, BD Biociences) and analyzed using Accrui 6, BD Bioscience software. At least 1×104 cells were collected and emission of the immunofluoresence of DCFH falls at FL2 channel i.e. at λ (wavelength) 530 nm (max). Oneparameter histograms based upon the florescence of the cell were generated via the analysis of the least 1×104 cells recorded.

### 8. Statistical Analysis

Significant variance between groups was performed for all groups using Student’s t-test. Data are expressed as means ± SD. Differences with P <0.05 were considered statistically significant.

## RESULTS

### 1. CCG promotes the monocyte-macrophage-like morphology of HL60 cells

The Giemsa stained control cells (treated with DMSO 0.1%v/v) exhibited undifferentiated HL60 cells which were round in shape and had unsegmented nuclei (fig. 1 right penal). In contrast to the control cells, the cells treated with crude extract of CCG exhibited strong adherence to plastic, a heterogeneous morphology characterized by increased cytoplasm to nuclear ratio, long cytoplasmic extensions, and multiple vesicles in their cytoplasm. The cells exhibited a spindle shape which is characteristic of macrophage-like cells (fig. 1 left penal). CCG does not affect the cell proliferation of HL-60 cells line as assessed by MTT assay (result is not shown).

**Figure 1.**
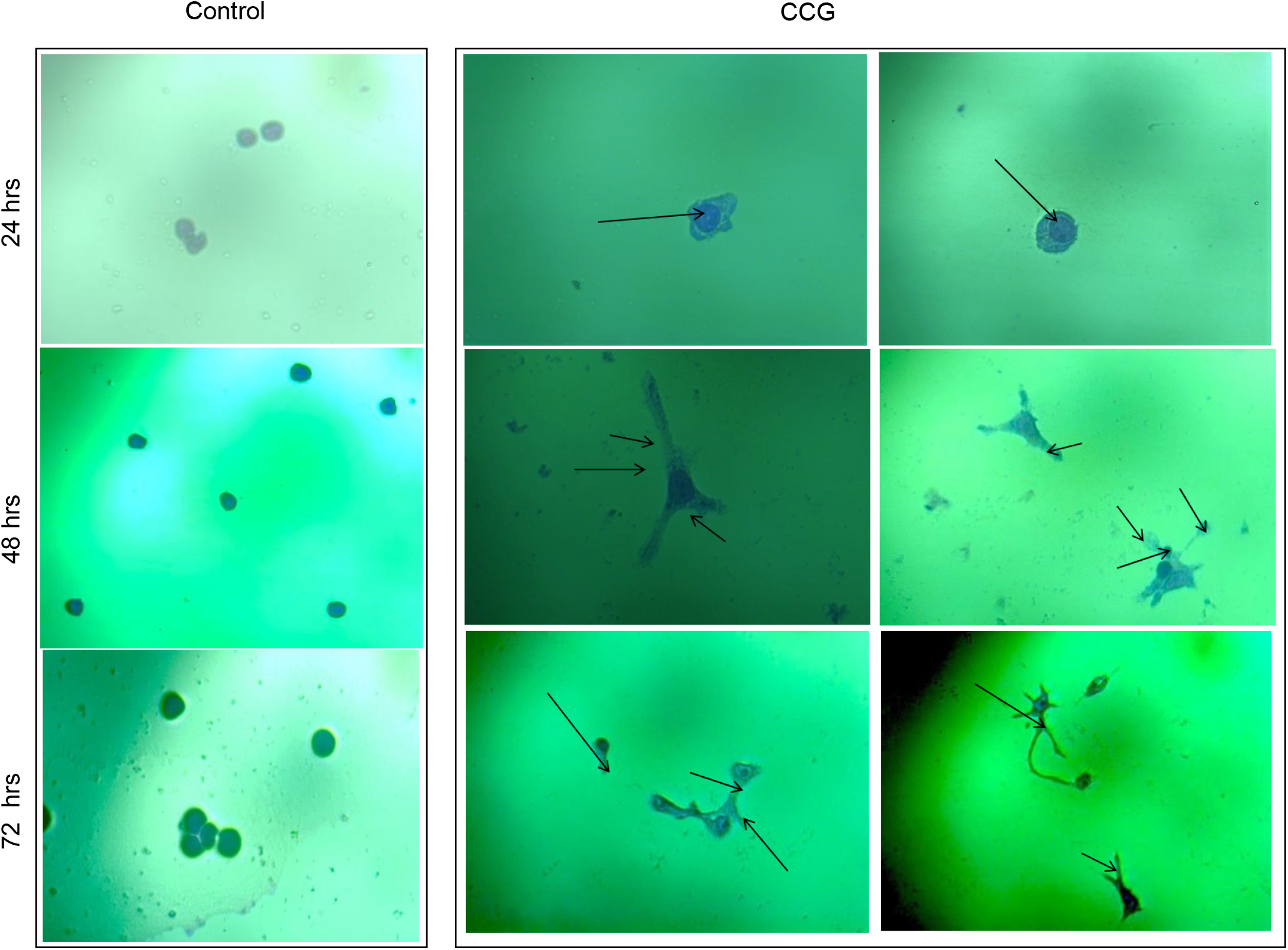
CCG promotes the monocyte-macrophage-like morphology of HL60 cells. HL60 cells were stained with Gemisa after treating with leaf crude extract of CCG for different time points. Control cells (Control) were treated with DMSO (0.1% V/V) Cells treated with 1µg/mL of leaf crude extract of CCG (CCG) at various time points and dose as mentioned. Arrow indicates the cytoplasmic extensions and vesicles in the cytoplasm. Images were captured in Leica phase contrast microscope 40X magnification. All experiments were performed at least three times.

### 2. CCG induces CD11b expression in HL60 cells

The crude extract of CCG induces the expression of CD11b in HL60 cells. The control cells(without any treatment) and cells treated with DMSO (0.1% v/v), used as vehicle, exhibit lowexpression of CD11b whereas treatment of crude extract of CCG (1μg/mL) for 24 hrs expression of CD11b is higher (fig. 2 A). The folds of increase of CD11b expression in crude extract treatment is ∼5.4 folds compared to the DMSO treated (fig. 2 B). CD11b is the Ill-subunit of the integrin cell surface receptor complement receptor 3 and functions in a heterodimer with CD18 to allow recognition and phagocytosis of iC3b opsonized particles. CD11b is exclusively expressed on the surface of mature monocytes, macrophages, neutrophils, and natural killer cells. Consequently, it is a widely used marker for monocyte/ macrophage differentiation (12). CCG also enhances the effect of PMA when treated together, which suggests that CCG might have a synergistic effect with PMA (figure is not shown).

**Figure 2:**
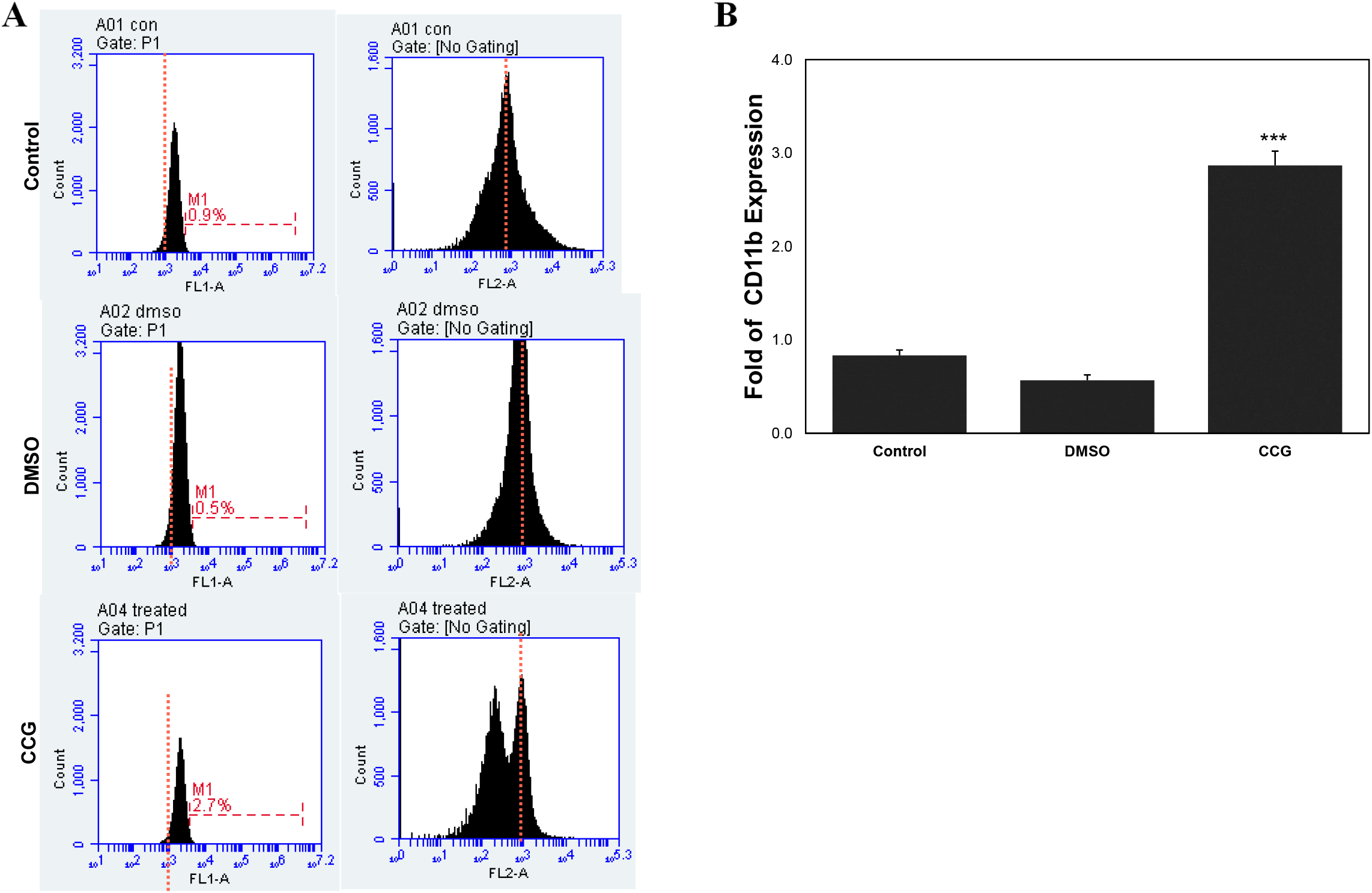
CCG induces CD11b expression in HL60 cells. (A) Histogram showing CD11b expression in control (Control), DMSO (DMSO) and treated cells with crude extract of CCG (CCG). (B) Graph showing fold increase of CD11b expression. Bar indicate standard deviation. *** indicates *P* <0.0001 compared to the control value. All experiments were performed at least three times.

### 3. CCG induces S phase cell cycle arrest

In order to determine the effect(s) of leaf crude extract of CCG on cell cycle progression, we subjected the control cells (treated with DMSO, 0.1% v/v) and treated cells with 1 ug/mL of leaf crude extract of CCG to flow cytometric analyses. Flow cytometry analyses of control cells indicated that almost 59.3, 31.2, and 8.6% of cells were distributed among G0/G1-, S-,and G2/M-phases, respectively. After treatment with leaf crude extract of CCG for 12 hrs remained comparable with that of control cells. However, the cells distribution of the control after 24 hrs among G0/G1, S, and G2/M-phases, were 38.2, 42.4, and 11.4% and for the treated the distribution were 23.1, 54.0, 9.6% respectively. Similarly, the cells distribution of the control after 96 hrs among G0/G1, S, and G2/M-phases, were 22.3, 62.1, 15.3%, and for the treated the distribution were 5.1, 77.2, 6.9% respectively (fig. 3 A). The results indicate that the percentage of cells in the G0/G1- and G2/M-phases decrease after treatment with crude extract whereas, the percentage of cells at S-phase increases after treatment in a time dependent manner (fig 3 B). A time-dependent increase in the early S-phase cell population along with a compensated decrease of cell in G0/G1- and G2/M-phases indicates that CCG induces cell cycle arrest at S-phase and cells are unable to proceed into G2/M-phase.CCG induces accumulation of ROS.The percentage of HL60 cells accumulating intercellular ROS was increased when cells were treated with CCG. The percentage of accumulating intercellular ROS in negative control cells (treated with DMSO, 0.1% v/v), positive control (treated with 100 mM of H_2_O_2_), and the cells treated with 1μg/mL of leaf crude extract of CCG were 7.2%, 26.4%, and 36.8%. respectively (fig 4 A). The fold of increase in the percentage of HL60 cells accumulating intercellular ROS is 3.67 in positive control whereas in the cells treated with leaf crudeextract of CCG it is 5.11 compared to the control cells (fig 4 B). The results indicate that the leaf crude extract of CCG has a very high ROS activity.

**Figure 3:**
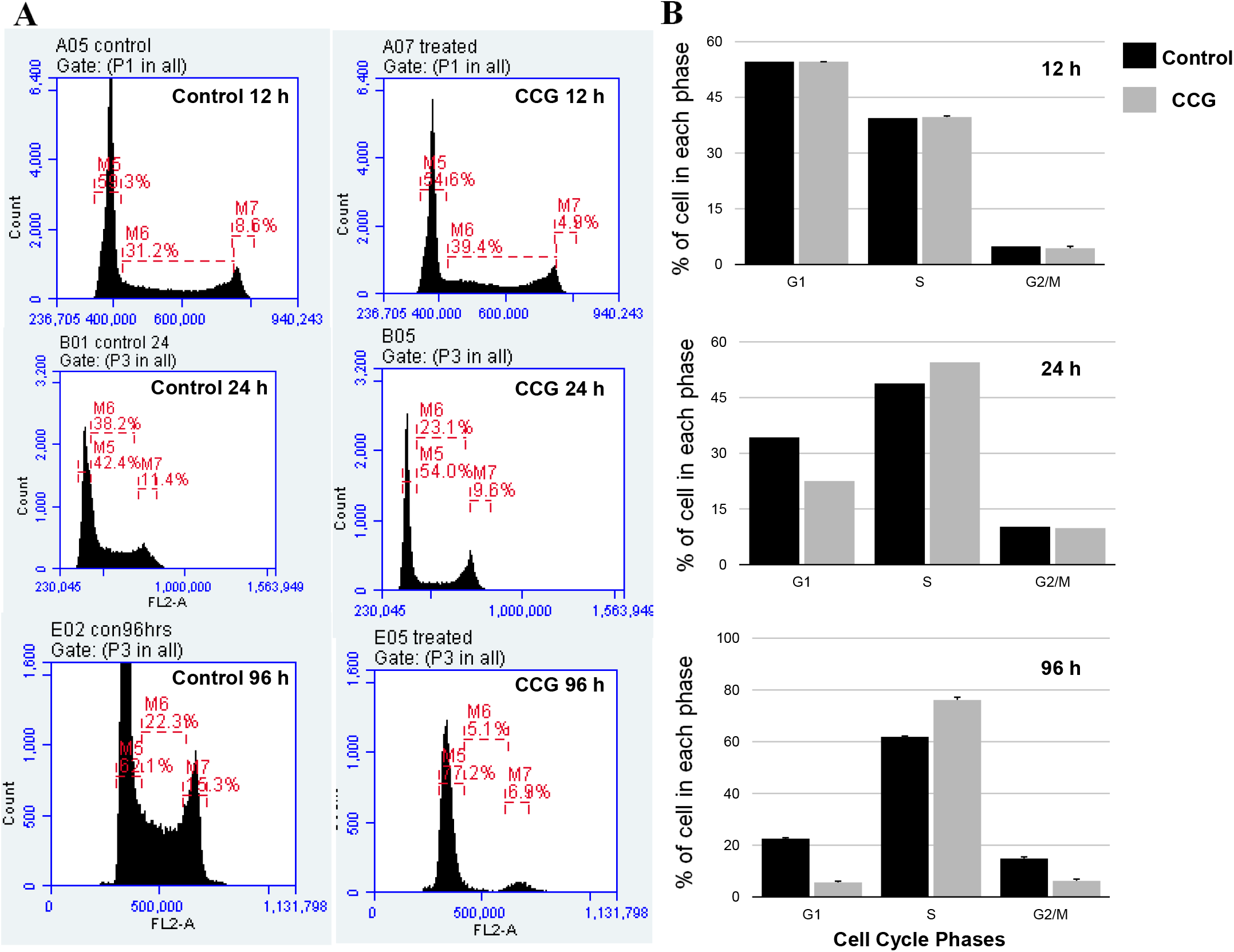
CCG induces S phase cell cycle arrest. Cell cycle was analyzed by flow cytometry for control (DMSO 0.1% v/v) and treated with leaf crude extract of CCG (CCG) for 12, 24 and 96 hrs. (A) Cell cycle histograms of cell cycle analysis at different time point are shown. (B) Graphs plotted for percentage of cells against cell cycle phases are shown. Bar indicates the error bar All experiments were performed at least three times.

**Figure 4.**
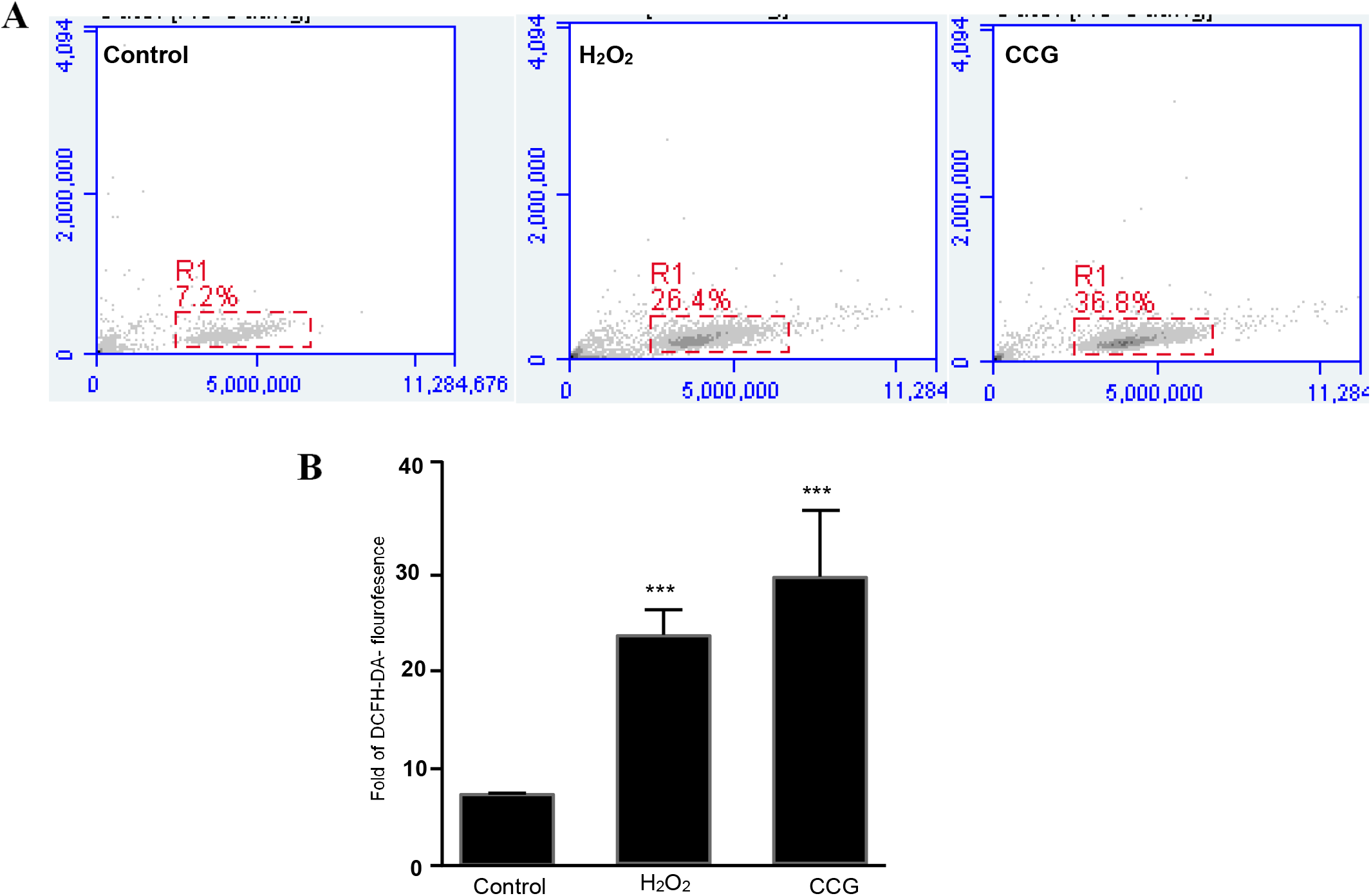
CCG induces accumulation of ROS. ROS generation was measured by flowcytometry as described in materials and methods. (A) Histograms of flowcytometry analysis of control, H2O2 and leaf crude extract of CCG (CCG) are shown. (D) Graph of DCFH-DA flouresence for control, H2O2 and leaf crude extract of CCG (CCG) was plotted. Bar indicates the error bar. *** indicates *P* <0.0001 compared to the control value. All experiments were performed at least three times.

## DISCISSION

Different extracts of *Croton caudatus* Geiseler (CCG) were practically non-toxic up to 150 mg/kg b. wt. and the LD50 was 350mg for aqueous and 500 mg and 650 mg for chloroform and ethanol extracts, respectively (13). CCG has been reported of having anticancer activity against human cancer cells derived from solid tumors. However, most of these reports are based only on the cytotoxicity effect or antioxidant activity of CCG. The present study investigates the anticancer effects of CCG on HL60. HL60, a human promyelocytic cell line, has been extensively used as an in vitro model for studying the effects of factors that regulate growth and differentiation of hematopoietic cells in general and of myeloid leukemia cells in particular. These cells proliferate as promyelocytes yet retain the capacity to undergo terminal myeloid or monocytic differentiation in response to various inducing agents. The results of the present study showed that crude leaf extract of CCG promotes the monocyte-macrophagelike differentiation of HL60 cells at low concentration (1μg/mL) as assessed by Giemsa staining although it does not affect the proliferation of HL60 cells as assessed by MTT assay. The observation of a heterogeneous morphology characterized by increased cytoplasm to nuclear ratio, long cytoplasmic extensions and multiple vesicles in their cytoplasm after treatment of leaf crude extract of CCG is similar to the cell differentiation into monocyte/macrophage (fig 1). The effect of leaf crude extract of CCG on the monocyte-macrophage like differentiation of HL60 cells was confirmed by the increased expression of surface CD11b which is the marker for monocyte/macrophage differentiation (fig 2).

The analysis of cell cycle of HL60 cells after treatment of leaf crude extract of CCG at different time points indicated that CCG induces cell cycle arrest at S phase (fig 3). Differentiating agents regulate the proliferation and myeloid maturation of HL60 cells by mechanisms that are at least partly independent (14). Our finding of CCG induces arrest of cell cycle at S phase is in accordance with the previous report that HL60 cells halted in G1 or S phase differentiate normally (15). The arrest of cell cycle at S phase may be due to the necessity of alteration in the expression of genes involved in HL60 differentiation since cells can launch differentiation while traversing S phase. Further, the results indicate that leaf crude extract of CCG has very high ROS activity as assessed by ROS accumulation detection using a flowcytometer (fig 4). ROS metabolism in AML may be targeted a therapeutic target.

## CONCLUSION

Since differentiation therapy is one of the promising strategies for the treatment of leukemia, universal effects have been focused on finding new differentiating agents. Therefore, the results of the present study showing that CCG indues HL60 cell differentiation and ROS accumulation without causing cytotoxicity to suggest CCG may be a promising differentiation induction chemotherapy (DIC) agent for the treatment of leukemia.

